# Chemotaxis and motility of *Achromatium oxaliferum* in response to oxygen, sulfide, and nitrate (*)

**DOI:** 10.64898/2026.03.30.715255

**Authors:** Sina Schorn, Danny Ionescu, Hans-Peter Grossart, Heribert Cypionka

**Affiliations:** Institute for Chemistry and Biology of the Marine Environment, Oldenburg, Germany; Max Planck Institute for Marine Microbiology, Bremen, Germany; Department of Environmental Microbiomics, Technische Universität Berlin, Berlin, Germany; Leibniz Institute of Freshwater Ecology and Inland Fisheries (IGB), Stechlin, Germany; Institute of Biochemistry and Biology, Potsdam University, Potsdam, Germany

**Author notes:** Department of Marine Sciences, University of Gothenburg, Gothenburg, Sweden (Sina Schorn). The authors have contributed equally to the work (Sina Schorn, Danny Ionescu). Dedicated to the memory of Hans-Dietrich Babenzien who has passed away, July 5, 2026, for his contribution to *Achromatium* research.

## Abstract

Bacteria of the genus *Achromatium* are known for their large cell sizes and intracellular calcium carbonate deposits. *Achromatium* cells inhabit freshwater, brackish, and marine sediments where they accumulate to high abundances at the oxic-anoxic interface. These bacteria alter their vertical position in the sediment along with daily fluctuations in oxygen concentrations. Yet, the mechanism behind their migration in the sediment remains unknown. In this study, we used chemotaxis assays and time-lapse microphotography to analyze the motility and chemotactic behavior of *Achromatium oxaliferum*. Microscopic observations revealed that rolling and gliding were the main forms of locomotion exhibited by *Achromatium*. In absence of any stimulant, the movement appeared to be mostly random and changes in direction frequently occurred. Chemotaxis assays showed a negative response of *Achromatium* to oxygen, sulfide, and nitrate, as evidenced by the change from undirected to directed locomotion against the respective chemical gradient. For periods of more than 1 hour, *Achromatium* cells moved continuously towards regions of low concentration. We further show that the genetic repertoire of *Achromatium* corresponds to our observations. Based on lab experiments and bioinformatic analyses we propose that *Achromatium* motility is propelled by type IV pili guided by a plethora of chemo- and photoreceptors. We further suggest that *Achromatium* uses negative chemo- and phototaxis to confine their distribution in aquatic sediments between opposing oxygen and sulfide gradients. This allows *Achromatium* to dynamically adjust its position in redox gradients, and thus is likely to have a major contribution to its success in the global colonization of diverse aquatic sediments.

## Introduction

*Achromatium* is the largest known freshwater bacterium but also inhabits brackish and marine sediments worldwide under a variety of environmental conditions [1–5]. It is known for its large intracellular deposits of (amorphous [6]) CaCO_3_ [1] and it’s hundreds of non-clonal chromosomes [7]. To date, there is no culture of *Achromatium* available and therefore little is known about the physiology and energy metabolism of these bacteria. It is assumed that *Achromatium* grows chemolithotrophically from the oxidation of inorganic sulfur linked to oxygen respiration [5, 8, 9], a hypothesis supported by genomic information. Environmental evidence for the proposed lifestyle was derived from the positive correlation between sulfate concentrations and the population size of *Achromatium* in a freshwater sediment [10], the intracellular deposition of sulfur globules, a typical feature of sulfur-oxidizing bacteria, and by the fact that *Achromatium* cells accumulate to high abundances at the oxygen-sulfide interface in the upper sediment layers [5]. Some sulfur bacteria, such as *Thiomargarita, Thioploca*, and *Beggiatoa* sp., use nitrate in addition to oxygen as an electron acceptor and they contain a large vacuole inside their cells to store it [11, 12]. This internal nitrate storage allows them to bridge the distance between oxygenated and sulfidic layers if the gradients do not spatially overlap. It remains unknown whether *Achromatium* is capable of reducing nitrate and couple it to sulfide oxidation as supporting experimental data [18] is contradicted by other experimental and genomic analyses [3, 10].

Experimental evidence indicates that some large sulfur-oxidizing bacteria use directed taxis to navigate chemical gradients of nitrate, oxygen, and sulfide in the environment. The filamentous sulfur bacterium *Thioploca* spp., for example, exhibits positive chemotaxis towards nitrate and low sulfide concentration, but exhibits a phobic response to oxygen and elevated sulfide [13]. A similar phobic response to elevated oxygen concentrations has been reported for *Beggiatoa* [14]. *Thiovulum* also exhibits chemotactic behavior to navigate to its preferred oxygen concentration and moving away when oxygen concentrations exceed this optimum [15]. Large magnetotactic bacteria likewise move along redox gradients, allowing them to scavenge and transport phosphate across the suboxic zone [16]. This behavior may be linked to the storage of energy in the form of polyphosphate, which they accumulate under favorable conditions and utilize when conditions become unfavorable [17].

*Achromatium* populations follow diurnal changes in the vertical depth of the oxic-anoxic interface by migration in the sediment, likely by a directed and active movement in response to oxygen availability [10]. It is, however, unknown if the vertical migration of *Achromatium* is controlled by chemotaxis to help the cells find their preferred zone in the sediment, as well as the mechanisms driving their vertical migration. We performed chemotaxis assays together with single cell tracking in artificial diffusion gradients to examine the motility and chemotactic behavior of *Achromatium* cells in response to oxygen, nitrate, and sulfide. We further describe the characteristic motility of *Achromatium* and their genetic repertoire for sensing chemical gradients and discuss the implications of our results for the lifestyle of *Achromatium* within opposing oxygen and sulfide concentration gradients in aquatic sediments.

## Results

### Motility pattern

*Achromatium* cells obtained from the upper layer of fresh sediments showed immediate locomotion upon contact with a glass surface. The presence of calcium carbonate bodies in the cells enabled them to sink passively to the surface within a minute of being suspended in water. We observed that cells whose calcium carbonate bodies had been experimentally removed often remained motile [19], but exhibited reduced motility, consisting of slight back and forth movements. The same type of movement was observed for cells that were naturally free of calcium carbonate bodies. However, the motility of calcium-free cells was not quantitatively assessed.

The movement of *Achromatium* cells was slow and often appeared jerky instead of continuous (Figure 1). Direct observation of the cells through a binocular or using video recordings revealed that most cells exhibited a “rolling” motion, which was characterized by the rotation of the cell along the short axis (Video S1). The rolling motion was sometimes interrupted by a flip over the long cell axis, during which the cells first moved into an upright position before dropping back down to the surface (Figure 1b). The cells remained in the upright position for a few seconds (Figure 1c, d). When cells flipped over the long cell axis, they often changed their direction of movement as well. We occasionally observed cells that exhibited a gliding motility, which proceeded along the long cell axis. Several cells were embedded in a mucus matrix; however, no mucus trail could be observed behind gliding cells, as typically seen in *Beggiatoa* spp. Exposure to blue light had an inhibitory effect on the motility of the cells which was reversed upon exposure to white or green light (Figure 1e).

**Figure 1.**
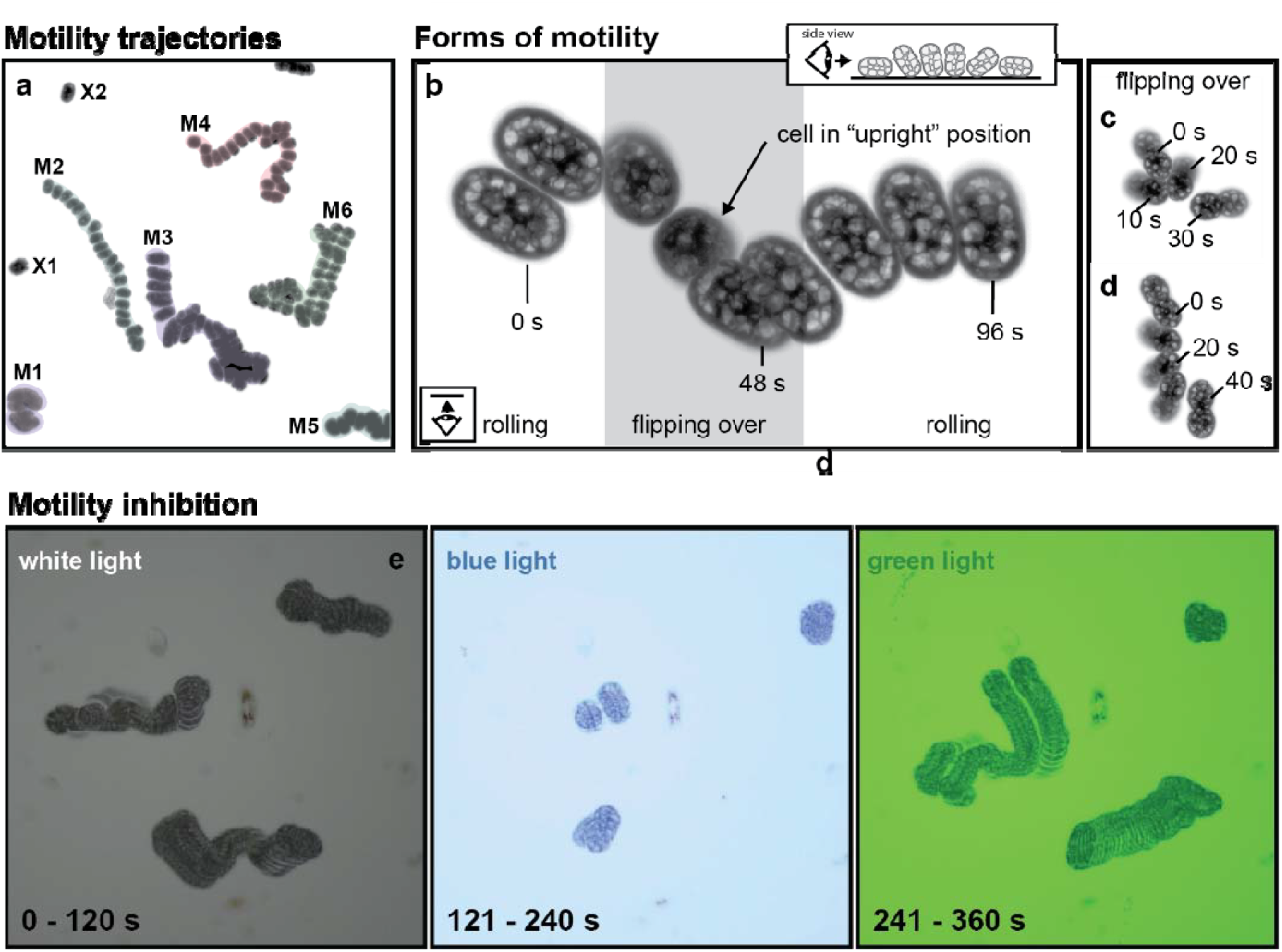
Motility characteristics of *Achromatium*. **a**. Typical trajectories of motile *Achromatium* cells on surfaces. The image is composed of 24 frames recorded every 10 seconds. Shown are trajectories of six motile *Achromatium* cells (M1-M6) as well as two cells which did not move (X1 and X2). **b**. Shown is the movement of a single cell over the duration of 96 seconds, represented as image stack of nine individual frames of a video recording. The inset shows the projected view of the cell if viewed from the side rather than from below. As illustrated in this example, most *Achromatium* cells moved by a lateral rolling motion of the cell body. The rolling motion was sometimes interrupted by a flipping over of the cell body, which could cause a change in direction. The flipping over movement can be seen as one end of the cell moved out of focus, resulting in the blurred structures visible in the images. **c, d**. While flipping over, the cells remained in an upright position for several seconds. In c, one cell is shown that remained in upright position for 30 seconds. In **d**., one cell is shown that moved into upright position, remained there for 20 seconds and then flipped over. Images were recorded with an inverted microscope, thus structures in focus were in contact with the surface of the microscope slide, whereas structures not in focus were in upright position. **e**. *Achromatium* movement was reversibly inhibited by exposure to blue light, i.e., when the cells were irradiated with dimmed white light, they showed normal movement (left panel), after applying a built-in blue filter (Zeiss Axioplan microscope) and an increased light intensity, movement was immediately stopped (middle panel). After switching to the green filter (same light intensity), cells continued their migration, with a few exceptions. Images were recorded every 2 seconds for a total of 120 seconds per light regime, meaning that every image is composed of 60 individual frames.

### Chemotactic behavior

Exposure of *Achromatium* cells to oxygen, sulfide, and nitrate resulted in a negative chemotactic response. This response was characterized by the transitioning from a random to a directional movement along a linear trajectory against the concentration gradient. The movement persisted for more than one hour (Figure 2 and Figure S1). No directional movement of *Achromatium* cells was observed in control experiments without a chemical gradient (Figure 2b). Upon exposure to oxygen, *Achromatium* cells migrated away from the oxygen source (air bubble) towards regions of lower concentration (Figure 2c). Exposure to sulfide also led to a negative response, with increasing concentrations (see methods) causing a more pronounced response and ultimately inhibiting cell motility (Figure S1). Conversely, no chemotactic response to thiosulfate was observed at the tested concentration. Nitrate and nitrite exposure produced less conclusive results; while some cells exhibited negative chemotaxis at lower concentrations, a clear negative chemotactic response was only observed at higher concentrations (>40 mM source; Figure S1). Overall, not all cells in the assay exhibited a chemotactic response; some cells were motile but moved randomly. The fraction of motile cells performing chemotaxis ranged from 10% to 51%, depending on the solvent and concentration tested (Table S1). Note that while the solute sources were at high concentrations, the diffusion into the aqueous chamber exposed the cells to environmentally relevant conditions for the duration of the experiments (see modeled diffusion in Fig. S1).

**Figure 2.**
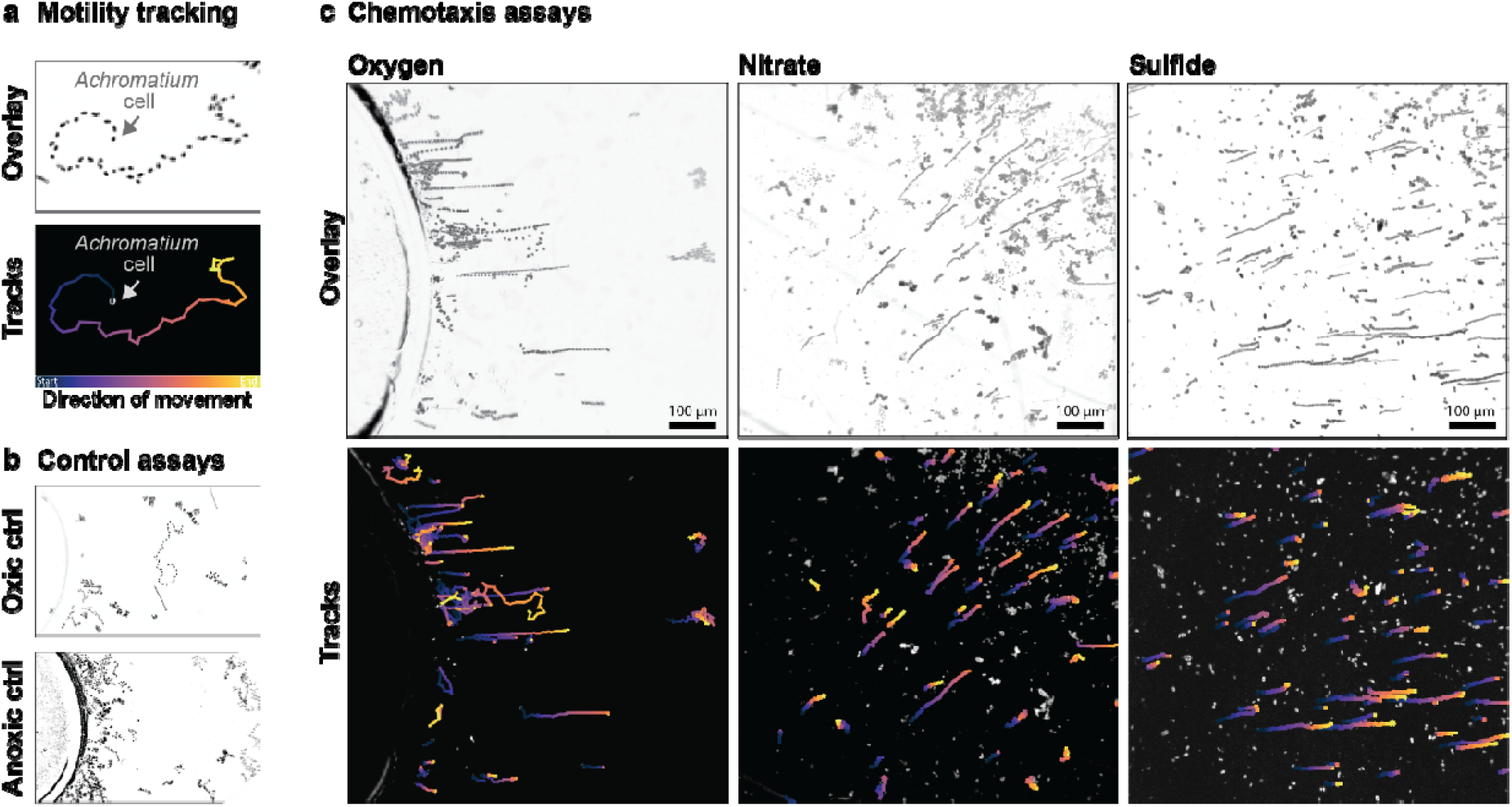
Chemotactic response of *Achromatium* cells to oxygen, sulfide, and nitrate concentration gradients. **a**. The movement of single *Achromatium* cells was followed by time lapse imaging and automated cell tracking. The tracks represent the trajectories of the cells, where the direction of movement is highlighted as a color gradient from blue to yellow. **b**. Control experiments were performed in which cells were either placed in oxic water and exposed to an air bubble (oxic control) or in anoxic water and exposed to a dinitrogen bubble (anoxic control). **c**. Chemotaxis assays showed the phobic response of *Achromatium* to oxygen, nitrate, and sulfide. *Achromatium* cells were placed in anoxic water and exposed to either an air bubble, sulfide-soaked agarose, or nitrate-soaked agarose. Note the different positions for the placement of the stimuli in the assays (lower left corner for nitrate and sulfide treatment). Overlays consist of 60 individual frames recorded every 30 seconds for 30 minutes (for oxygen) or 60 seconds over a period of 1 hour (for nitrate and sulfide).

### Single cell velocities

We compared the velocities of cells moving in chemical gradients with those of cells moving in media without a gradient (Fig. 3). Median velocities were 1.57 µm s^−1^ in the oxygen gradient, 0.38 µm s^−1^ in the nitrate gradient, and 0.37 µm s^−1^ in the sulfide gradient. Under uniform anoxic, oxic (94% O_2_), and hypoxic (5% O_2_) conditions, median velocities were 0.44, 0.91, and 2.07 µm s^−1^, respectively.

**Figure 3.**
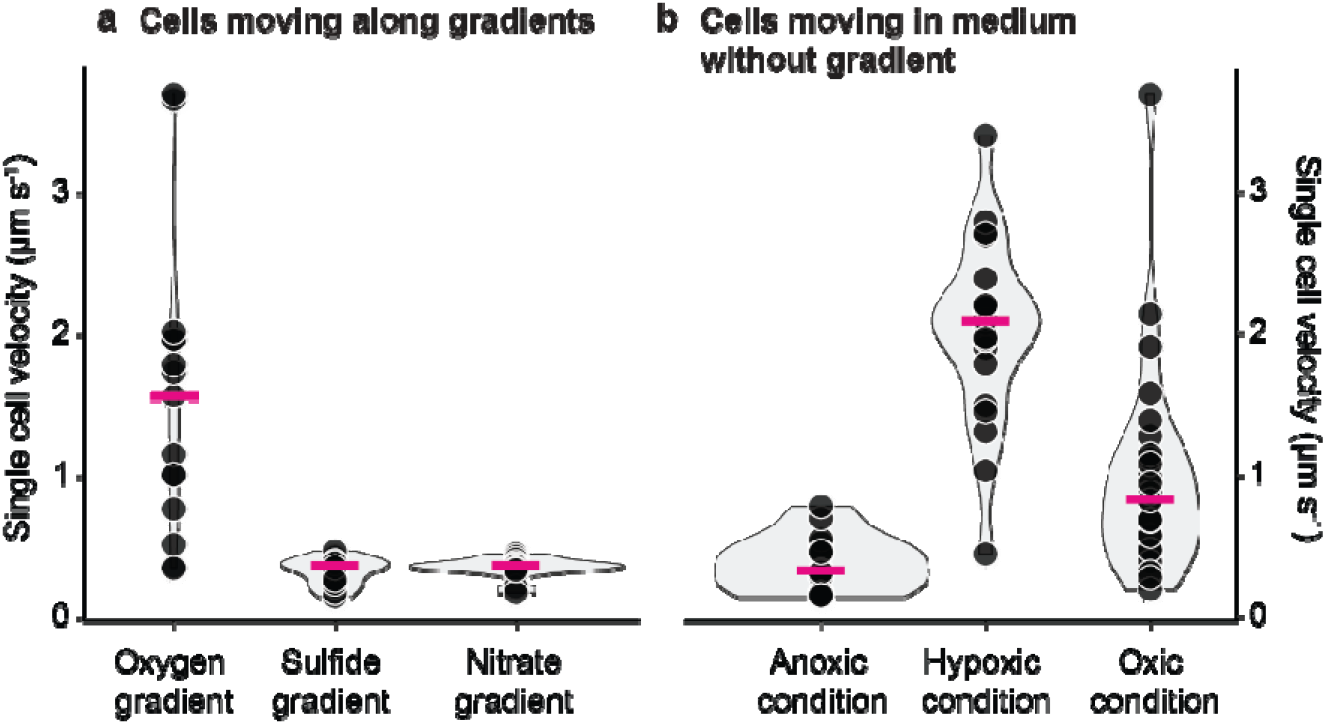
Comparison of single cell velocities. Single cell velocities were determined for periods of continuous movement, either for cells exposed to chemical gradients of oxygen, sulfide, and nitrate (**a**) or under uniform oxic (94% O_2_), hypoxic (5% O_2_) and anoxic conditions (**b**). Each dot in the plot represents the velocity of a single *Achromatium* cell, the density (width) and the distribution (height) of the data is represented by the shape of the violin plot.

Velocity distributions differed significantly among the six conditions (Kruskal–Wallis H(5) = 72.09, p = 3.76 × 10^−14^). Dunn’s post hoc tests showed that velocities in the oxygen gradient were significantl higher than those in the nitrate gradient (Bonferroni-adjusted p = 1.26 × 10^−4^), sulfide gradient (p = 6.43 × 10^−5^), and anoxic control (p = 0.0358). Velocities under oxic conditions were significantly higher than those in the nitrate (p = 4.54 × 10^−4^) and sulfide gradients (p = 2.25 × 10^−4^). Velocities under hypoxi conditions were significantly higher than those in the nitrate gradient (p = 1.97 × 10^−9^), sulfide gradient (p = 9.39 × 10^−10^), and anoxic control (p = 1.67 × 10^−5^). No other pairwise comparisons were significant after correction (adjusted p ≥ 0.0668).

The directionality of cell movement was quantified using the chemotaxis angle and a signed chemotaxis index [20, 21]. The chemotaxis angle was defined relative to the direction toward the chemical source, such that 0° indicated movement directly toward the source, 90° movement parallel to the source boundary, and 180° movement directly away from the source. The estimated median chemotaxis angles were approximately 175° for oxygen, 166° for nitrate and 148° for sulfide, corresponding to deviations of approximately 5°, 14° and 32°, respectively, from movement directly away from the source. The median chemotaxis indices were −0.68 (IQR: −0.79 to −0.38) for oxygen, −0.69 (IQR: −0.86 to −0.40) for nitrate and −0.54 (IQR: −0.68 to −0.36) for sulfide. The corresponding mean ± SD values were −0.55 ± 0.33, −0.59 ± 0.34 and −0.51 ± 0.24, respectively. The predominantly negative values demonstrate net movement away from all three chemical sources, although a small number of individual trajectories showed neutral or weak movement toward the source.

### Chemotaxis-related genes

Protein annotation of *Achromatium* genomes was already conducted and detailed in our previous studies [3, 7]. Genome resolved transcriptomic [3] and proteomic [22] data from parallel studies provided complementary evidence for expression of a type IV pilus system in *Achromatium*. Transcripts were detected for structural pilins and proteins involved in pilin processing, pilus assembly, extension and retraction, including PilB, PilC, PilD, PilM, PilN, PilO, PilP and PilT. Proteomic analysis additionally detected major and minor pilins (PilA/PilE), the outer-membrane secretin PilQ, the extension ATPase PilB, the retraction ATPases PilT and PilU, and several accessory and regulatory proteins, including PilV, PilW, PilX, PilY1 and PilGHIJL. Together, the two datasets cover the principal components required for pilin processing, filament formation, trans-envelope assembly, extension and retraction (Fig. 4).

In the present study, we focused on the identification of chemotaxis receptors. Our analysis revealed that *Achromatium* has multiple mechanisms for signal-dependent motility, i.e. chemo- and phototaxis (Fig. 4). First, it harbors multiple canonical methyl-accepting chemotaxis proteins (MCPs) with a variety of sensing domains, including PAS for redox and oxygen sensing, TarH / 4HB (4-helix bundle) for amino acids and small organics, Cache (sCache and dCache) for amino acids, polyamines, organic acids, CHASE for peptides and hormones, and BLUF (blue-light photoreceptor domain that uses a flavin cofactor (FAD) to sense light) domains for blue-light sensing (Table S2; [23]). Second, it has a nitrate-binding domain of pilJ [24]. Additionally, it harbors several blue-light regulated diguanylate cyclases which regulate the abundance of cyclic-di-GMP and are known to control phototaxis and switching from motile to sessile states [25] (Table S3). Last, *Achromatium* also harbors several complex two-component system light/redox sensors in the form of PYP-PAS sensory histidine kinases (Table S3). Sequences of all identified proteins are provided in the supplementary material. The sequences and domain characterization (extracted from InterScanPro annotation) are available on the Zenodo repository at: https://doi.org/10.64898/2026.03.30.715255.

**Figure. 4.**
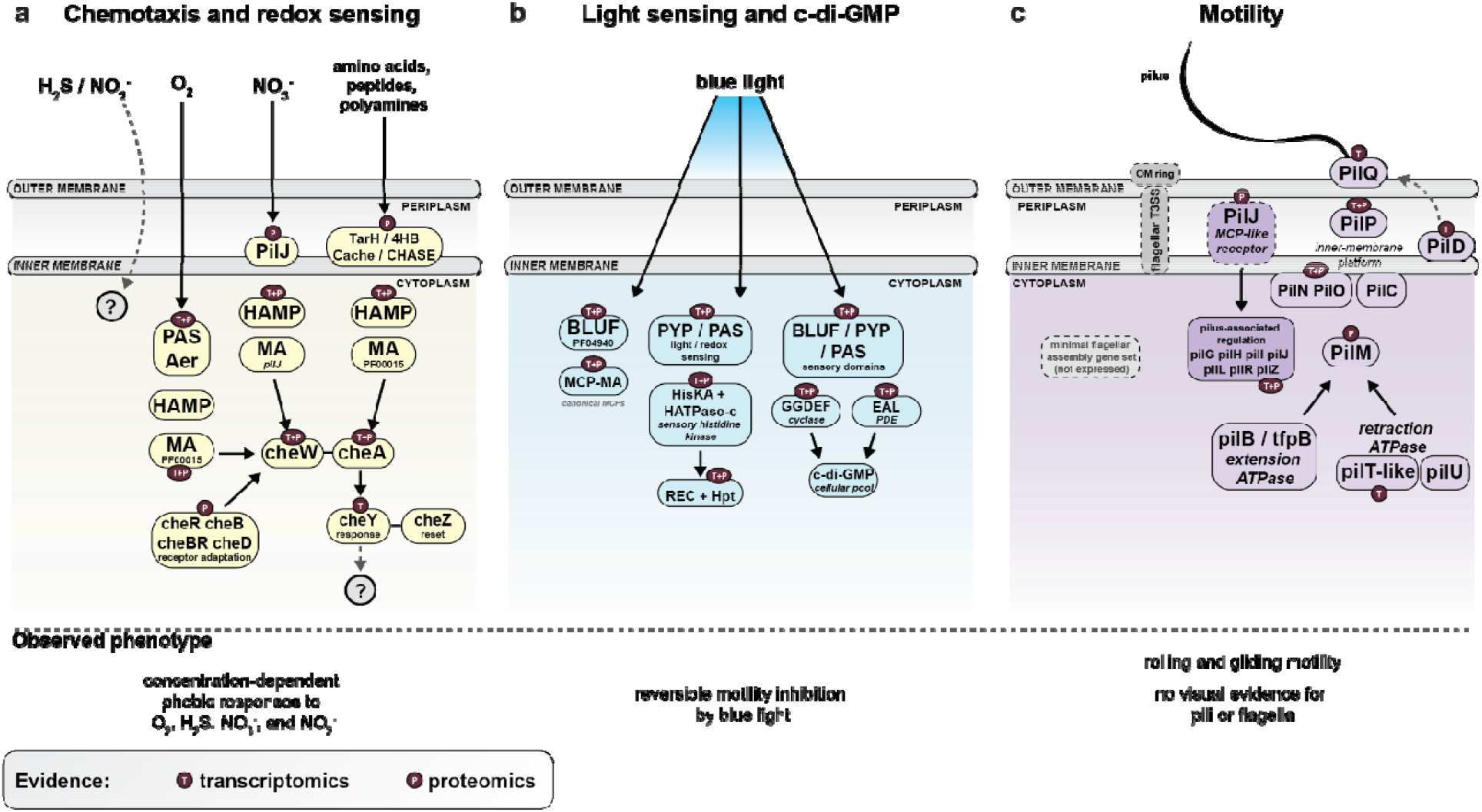
Genomic and transcriptomic evidence for sensory and motility systems in *Achromatium*. a.Representative methyl-accepting chemotaxis proteins (MCPs) contain PAS/Aer-type domains associated with intracellular redox or oxygen sensing, PilJ-type domains potentially involved in nitrate sensing, and TarH/4HB, Cache and CHASE domains predicted to recognize amino acids, organic compounds, polyamines or peptides. Chemotactic responses to sulfide and nitrite were observed experimentally, but the corresponding receptors remain unidentified. MCP signals may be transmitted through HAMP and methyl-accepting (MA) domains to the Che signaling system. *Achromatium* genomes encode *cheA, cheB, cheBR, cheD, cheR, cheW, cheY* and *cheZ; transcripts of cheA, cheB, cheR, cheW* and *cheY* were detected in the Lake Stechlin transcriptome. **b**. Identified light-sensing systems include BLUF-containing MCPs, PYP/PAS-containing sensory histidine kinases and proteins combining BLUF, PYP or PAS domains with GGDEF and/or EAL domains. GGDEF and EAL domains are predicted to regulate the cellular c-di-GMP pool, potentially linking light perception to the observed reversible inhibition of motility by blue light. The intracellular localization and regulatory connections of these proteins have not been experimentally resolved. **c**. *Achromatium* genomes encode the type IV pilus-associated proteins PilG, PilH, PilI, PilJ, PilT and PilU, consistent with the proposed involvement of pili in the observed rolling or twitching-like movement. Gliding was also observed, although its mechanism remains unresolved and may involve the mucus envelope. A minimal flagellar assembly/type III secretion system is present in the genomes, but no flagellar filament has been observed and flagellar genes were not expressed in the examined Lake Stechlin transcriptome. Dashed arrows indicate proposed or unresolved regulatory connections rather than experimentally demonstrated interactions. Domain architectures and membrane positions are simplified.

## Discussion

### Negative chemotactic responses may help to confine the distribution of *Achromatium* in sediments

The negative chemotactic response of *Achromatium* to oxygen and sulfide observed in our assays suggests that *Achromatium* migrates vertically towards regions of low oxygen and low sulfide concentrations. A similar chemotactic response to oxygen and sulfide is known for other sulfur-oxidizing bacteria, e.g. *Beggiatoa* and *Thioploca* [13, 15, 26, 27]. The chemotactic response of *Thioploca* and *Beggiatoa* to sulfide was shown to be concentration-dependent and was negative at elevated concentrations but positive at lower, micromolar concentrations [13]. *Achromatium* also revealed a concentration-dependent response, with a higher number of directionally motile cells at low and intermediate sulfide concentration (1 mM and 5 mM source; Fig. S1), and only random movement at higher (20 mM source) concentrations (Fig S1).

Furthermore, *Achromatium* displayed negative chemotaxis to nitrate. *Achromatium* cells are often found in anoxic sediments as well and were hypothesized to couple nitrate reduction to sulfide oxidation [18]. However, there is so far no conclusive experimental [10] or genomic [3] evidence for denitrification activity or for the capacity to store nitrate in *Achromatium* cells. This is in contrast to other large sulfur-oxidizing bacteria, such as *Beggiatoa, Thioploca*, and *Thiomargarita*, that maintain intracellular reservoirs of nitrates to be used as an electron acceptor. Accordingly, *Thioploca* responds positively towards nitrate [13]. This storage capacity allows them to bridge the distances between the oxygenated and sulfide layers when oxygen and sulfide gradients do not spatially overlap. Our results confirm previous genetic analyses of *Achromatium* [3] suggesting that its cells don’t use nitrate for anaerobic respiration as they lack genes encoding the respiratory nitrate reductases (Nar). Nevertheless, *Achromatium* genomes encode a nitrite reductase (NirK) and nitric oxide reductase (NorBC), indicating the potential to reduce nitrite to nitrous oxide.

*Achromatium* displays nitrate-driven negative chemotaxis only when exposed to the highest concentrations used in this experiment (source of 50 and 100 mM; Fig. S1). In comparison, negative chemotaxis in response to nitrite was observed at all tested concentrations (source of 1-100 mM; Fig. S1). In aquatic sediments, nitrate often extends below the oxygen penetration depth and nitrite, even though it may not accumulate, may be dynamically formed and consumed. Thus, nitrate and nitrite could act as another oxidizing cue once oxygen is depleted and signal the cells to venture deeper into the anoxic zone until sulfide starts to accumulate. Alternatively, evasion of high nitrate concentrations may be a mechanism to prevent accumulation of nitrite in the cells. *Achromatium* may not be capable of denitrification or DNRA, yet it possesses periplasmatic nitrate reductase (PF02613/65) whose activity may result in the accumulation of nitrite. And, while it appears *Achromatium* also possesses a nitrite reductase, rate imbalances have been shown in other bacteria to lead to nitrite accumulation [28]. Inspection of transcriptomic data of *Achromatium* [3] showed that both nitrate and nitrite reductase were transcribed whereby the nitrate reductase transcription was 10-fold higher than that of the nitrite reductase. The latter difference in expression levels supports the potential accumulation of nitrite due to activity imbalance between enzymes.

The chemotaxis indices demonstrate that *Achromatium* cells maintained a substantial directional movement relative to the stimulus source. The absolute median indices of approximately 0.54–0.69 indicate that more than half of the distance travelled contributed to net displacement along the stimulus axis. Direct comparisons with other bacteria are limited. Bacterial chemotaxis studies generally report a population-level relative chemotactic velocity, (V_chemotaxis_/V), rather than the median of trajectory-specific chemotaxis indices. Nevertheless, (V_chemotaxis_ /V) has the same underlying interpretation as a path-length-weighted chemotaxis index, i.e. the fraction of movement contributing to net displacement along the gradient. Reported values for flagellated bacteria include 0.06 and 0.14 for *Escherichia coli* in aspartate and serine gradients, respectively, 0.23 for *Salmonella Typhimurium* in a serine gradient, and 0.018–0.35 for *E. coli* exposed to different α-methylaspartate gradients [29–31]. For the marine flagellated bacterium Vibrio *alginolyticus*, less than 10% of the swimming velocity was directed up a serine gradient [32].

The ability of the slowly moving *Achromatium* cells to maintain gradient-aligned trajectories is notable. Their large dimensions could increase the concentration difference occurring across the cell and reduce Brownian reorientation. However, active rolling complicates a simple model based on comparison between fixed receptor regions. If cell displacement occurs perpendicular to the long axis, balancing the signals received at the two ends could orient the long axis along an isoconcentration line and thereby direct lateral movement along the gradient. Alternatively, rolling could allow localized receptor regions to scan the gradient and generate phase-dependent changes in pilus activity, resembling klinotaxis. A combination of spatial and temporal sensing is also possible. Short-axis rotation was previously described in a considerably smaller bipolar bacterium, for which both a two-pole spatial-sensing model and rotation-assisted klinotaxis were considered [33]. Direct spatial sensing has also been demonstrated in non-rolling, surface-attached *Pseudomonas aeruginosa* [34]. Whether *Achromatium* uses simultaneous spatial comparison, rotation-assisted scanning, temporal sensing, or a combination of these mechanisms remains unresolved.

### Genomic potential for chemotaxis and motility of *Achromatium*

Due to its genomic complexity and intracellular heterozygosity [3, 7] no closed *Achromatium* genome has been obtained so far. Nevertheless, several high completeness single-cell and metagenome assembled genomes of *Achromatium* from freshwater [3, 7] and saline environments [4, 35] are available. These allowed us to search for chemotaxis-related genes, some of which are also depicted in Ionescu et al. [3].

*Achromatium’s* motility is surface-dependent and is likely propelled by pili. *Achromatium* spp. from geographically distant locations possess key genes involved in motility [3, 4], including the full set of genes encoding for type IV pili (*pilGHIJTU*) for twitching motility. Phenotypically, the observed rolling motility of *Achromatium* is reminiscent of twitching motility as described in the literature, which is the surface-associated movement facilitated by type IV pili [36]. The overall velocity exhibited by *Achromatium* cells (<2.3 µm s^−1^) is comparable to other twitching bacteria, such as *Pseudomonas aeruginosa* (< 0.2 μm s^−1^) [37] or *Neisseria* species (ca. 1.6 µm s^−1^) [38]. *Achromatium* shares other motility patterns with twitching bacteria, including the ability to stand upright on a surface. This upright positioning has been previously described for P. aeruginosa too and is believed to facilitate 3D-sensing of the environment [39]. We therefore propose that pili-mediated twitching underlies the rolling locomotion of *Achromatium*. Although there is so far no conclusive visual evidence for the presence of type IV pili by *Achromatium*, these are hard to identify in TEM or SEM images when not directly targeted.

*Achromatium* additionally exhibited gliding motility. Both forms of motility, rolling and gliding, are common among large sulfur bacteria [27]. Gliding motility in filamentous sulfur bacteria was suggested to results from propulsion of the cell body through excretion of mucus through pores on the cell surface [41], similar to gliding cyanobacterial filaments [42]. *Achromatium* cells also exhibit mucus envelopes [19, 40], therefore, a similar mechanism of propulsion may underlie the observed gliding motility.

*Achromatium*, possesses the minimal set of genes necessary for flagellar assembly as identified in Liu et al. [43, 44]. Additionally, it possess a complete set of the chemotaxis signaling genes *cheA, cheB, cheBR, cheD, cheR, cheW, cheY, cheZ*, which are known to regulate the movement of the bacterial flagella. Nevertheless, in over a century of microscopic investigations of *Achromatium*, no flagella were ever reported. While we cannot rule out a conditional expression of a flagellum, it is likely that the chemotaxis machinery has been rewired toward a pili-based system, as mentioned above. Similar architectures are known in other bacteria. In Pseudomonas aeruginosa, a dedicated Pil-Chp chemosensory pathway with the MCP-like receptor PilJ and Che-like regulators PilG/PilH controls Type IV pili chemotaxis and mechanotaxis rather than flagellar rotation, demonstrating direct coupling of Che-type signaling to Type IV pili taxis [45, 46]. In the non-flagellated bacterium, *Lysobacter enzymogenes*, homologous components of the flagellar type III secretion apparatus have been experimentally shown to acquire a novel function in controlling twitching motility, indicating that flagellar export/basal body modules can be repurposed to regulate T4P dynamics in the absence of any filament [47].

Investigating the transcriptomes of *Achromatium* from Lake Stechlin [3], revealed that under the conditions prevailing during sampling, no flagellar genes were expressed, in contrast to all Type IV pili genes. Additionally, we could confirm the expression of the Type IV pili MCP, *pilJ* and of the chemotaxis signaling genes *cheA, cheB, cheR, cheW, and cheY*. Thus, it seems likely that *Achromatium* performs a pili driven motility guided by the che signaling proteins. A parallel proteomics study [22] confirmed the expression of Type IV pili associated genes, so that together with the previous transcriptomic data, the expression of all necessary genes can be confirmed.

*Achromatium’s* response to blue light is also strongly supported by genomic evidence. First, we recovered several canonical MCPs containing a BLUF domain. Second, we identified multiple light sensing proteins with both PYP and BLUF domains coupled to the regulation of GGDEF (diguanylate cyclase), EAL (phosphodiesterases), or both in dual GGDEF/EAL motifs. GGDEF leads to the production and accumulation of cyclic-di-GMP, while EAL degrades it. Together, these activities are known to facilitate the switch between sessile and motile states of cells [48–50]. In *Achromatium*, we propose that blue light results in an increase of c-di-GMP and no motility, whereas removal of blue light or exposure to other light reduces c-di-GMP concentrations and allows for motility.

Finally, *Achromatium* also harbors complex two-component system light/redox sensors in the form of PYP-PAS sensory histidine kinases. These were detected with several levels of complexity allowing for fast on/off responses as well as improved regulation[51]. Previously, such systems have been shown to be involved in the regulation of pili-based motility in other bacteria [52, 53].

Based on the data at hand, we propose that *Achromatium* cells may have abandoned flagellar locomotion, in favor of Type IV based twitching motility. At the moment, it remains unclear whether *Achromatium* maintains the ability to conditionally express flagella or whether it has repurposed the Type 3 secretion system, which forms the basis of the flagellar mechanism [54]. The presence of such a variety of chemoreceptors and motility regulation systems harbored by *Achromatium* support the notion that regulation of the pili-based motility is far more complex than that of the flagella based one [55].

The high density of *Achromatium* cells resulting from their CaCO_3_ inclusions is unlikely to preclude type IV pilus-mediated motility. Type IV pili are powerful molecular motors: individual pili can generate retraction forces of approximately 50–100 pN [56], while coordinated pilus bundles can reach the nanonewton range [57]. Moreover, the accessory retraction ATPase PilU can support type IV pilus activity under high-friction conditions [58]. At the observed cell dimensions and velocities, *Achromatium* moves at a Reynolds number far below one (Re ~10^−5^ – 10^−4^ for cells 50-100 µm long in water); inertia is therefore negligible, and hydrodynamic drag alone is likely small relative to the force generated by a single retracting pilus. Yet it remains unknown how a type IV pili-mediated locomotion system operates in such unusually large, mineral-loaded cells.

## Conclusion

Our chemotaxis assays reveal that through negative chemotaxis *Achromatium* is able to position itself in the sediment in its preferred niche, i.e. at the sulfide-oxygen interface. For its precise positioning in the sediment, negative chemotaxis, i.e. evading high concentrations of, e.g., oxygen or sulfide, allows the cell to simultaneously respond to concentration changes of multiple solutes, which is more favorable than movement towards higher concentrations of just a single solute. Single-cell velocities of *Achromatium* are slow compared with those of other chemotactic cells (<2 µm s^−1^ versus >10 µm s^−1^) [59, 60], limiting their ability to respond rapidly to changing environmental conditions. Instead, persistent but slow movement away from unfavorable conditions may enable cells to eventually locate favorable conditions at the oxygen–sulfide interface and follow its vertical shifts over the day–night cycle.

Additionally, *Achromatium* appears to be well equipped to perceive blue light. This likely compensates for its ability to sense redox gradients and allows it to respond to day/night cycles which in turn can drive diurnal migration of the sulfide-oxygen interface due to the interplay of photosynthesis and respiration. This response likely explains the reported change in *Achromatium* cells distribution during illuminated and dark periods [10]. *Achromatium* has been suggested to survive in many different environments [3]. It is likely that its complex systems of perceiving its chemical and photic environment, are part of the cell’s ability of a fine-tuned response to various environmental changes, not only through expressing various alleles of the same gene as suggested [3, 7], but also through accurate positioning in the sediment, for optimal activity. Thus chemotaxis is likely a major contributor to *Achromatium’s* suggested ubiquitous presence [3] in aquatic sediments where it can sense and benefit from redox gradients.

## Material and Methods

### Sampling and storage of *Achromatium* cells

*Achromatium* cells were collected from the sediments of Lake Stechlin located in Northeastern Germany, Brandenburg (Germany), at the pier of the Leibniz Institute of Freshwater Ecology and Inland Fisheries. Surface sediment with fresh *Achromatium* cells was retrieved with beakers from the upper 3 cm of the lake sediment and filled into jars to about a height of 3 cm, and topped with ca. 5 cm of lake water. The jars were stored at 15 °C at a 12 h/12 h light/dark cycle.

### Collection and purification of *Achromatium* cells

*Achromatium* cells were always freshly harvested from the jars prior to experiments and microscopic observations. To separate *Achromatium* cells from large sand grains, the sediment was filtered through an 80-µm pore size mesh. The collected cells were then subjected to gentle rotation in a glass petri dish to separate them further from organic debris and sediment particles. The resulting purified *Achromatium* cells were transferred to a new dish containing sterilized, filtered lake water, and the rotation steps were repeated until a visibly clean population was obtained. This protocol was adapted from previous studies [5].

### Chemotaxis assays

The *Achromatium* cells used for the chemotaxis assays were purified under anoxic conditions in an anaerobic chamber filled with an N_2_/CO_2_ gas and suspended in sterile-filtered, anoxic lake water. The purified cells were then placed in a horizontal chemotaxis chamber consisting of a microscope slide and frame with an opening for the introduction of a chemical stimulus. The chamber was 16.5 × 16.5 × 2 mm L×W×H with the solute opening being 2 mm wide. To prevent oxygen diffusion during microscopic observation, the chamber was sealed with a glass coverslip and paraffin oil on the edges. To test the chemotactic behavior of *Achromatium*, we exposed the cells to a range of stimuli including oxygen, sulfide, nitrate, nitrite, and thiosulfate. Oxygen was introduced via an air bubble at the chamber opening to promote diffusion of oxygen into the water. For sulfide, nitrate, nitrite, and thiosulfate, we employed an agarose block soaked to saturation (>12 h) with solutions of each compound at varying concentrations ranging from 1 to 100 mM, as detailed in Table S1. Two control experiments were conducted to assess the non-chemotactic behavior of *Achromatium* cells under different oxygen conditions. In the first control, cells were exposed to an air bubble while being in fully oxygenated water (oxic control). In the second control, conducted in an anoxic tent, cells were exposed to a dinitrogen bubble while being in anoxic water (anoxic control). The motility pattern of the cells was then followed microscopically by recording images at regular intervals of 30 to 60 s over a 30 to 60-minute period. The trajectory of individual cells was reconstructed by overlaying the individual frames, as illustrated in Figure 1b.

### Light exposure experiments

Light experiments were conducted using a Zeiss Axioplan microscope equipped with built-in filter cubes for blue and green light using the same light intensity for both the blue- and green-light exposures. Repetitions of the experiment resulted in a similar response. Movement of *Achromatium* cells in response to white, blue, and green light was recorded every 2 seconds for a total of 120 seconds per light regime.

### Motility pattern of *Achromatium*

We observed the motility of *Achromatium* cells on glass slides after purification from the sediment. These observations were used to characterize the different forms of motility. Our observations included cells containing calcium carbonate bodies, as well as cells that were naturally devoid of calcium carbonate or from which the calcium carbonate bodies had been experimentally removed using dilute acids, ethanol, ethylenediaminetetraacetic acid (EDTA), sodium bicarbonate, or UV radiation as described in Schorn et al [19].

### Motility pattern of *Achromatium* under different oxygen concentrations

To investigate how *Achromatium* cells respond to variations in oxygen concentration, with regard to changes in velocity, we utilized cells obtained from sediments collected in 2016 and September 2022. The sediment was stored in jars at room temperature (approximately 22 °C) under standard light conditions. To prevent cell damage due to the weight of the coverslip during microscopic observation, we used electrical tape to create a frame on the slide, high enough to allow the cells to move freely. *Achromatium* cells were either placed into oxygenated water or water degassed with N_2_. We refer to these conditions as oxic and anoxic. We repeated these experiments in 2022, this time measuring the oxygen content in the water during the observation. We performed two experiments, one containing 94% O_2_ saturation at 21.9 °C and one containing 5% O_2_ saturation at 21.9°C. We refer to these conditions as oxic and hypoxic. The oxygen content in the water was measured with a needle optode (Pyroscience OXR430). The motility of *Achromatium* cells was recorded on an inverted microscope (Leica DMI 6000B) by taking images from the same field of view every 10 to 30 seconds over a time period of several minutes.

### Chemotaxis indices

Chemotaxis angles and indices were estimated from the color-coded trajectory overlays shown in Fig. 2 using a custom Python script (Python 3.12.14; NumPy 2.3.5, SciPy 1.17.0, Pillow 12.3.0 and Matplotlib 3.10.8). Colored trajectory pixels were separated from the greyscale images in HSV color space, and individual trajectories were identified as connected components. Because track color encoded time, the temporal order of each trajectory was reconstructed from its hue, and a centerline was generated from the centroids of 12 successive color intervals. Components containing fewer than 22 colored pixels, spanning less than 43% of the color-time scale, containing fewer than six valid intervals or having a reconstructed path shorter than for pixels were excluded. For each retained trajectory, net displacement was calculated between the reconstructed start and end points, while total path length was calculated as the sum of the successive centerline displacements. The direction toward the chemical source was defined as perpendicular to the local bubble boundary for oxygen and toward the lower-left side of the diagonal source boundary for nitrate and sulfide. The chemotaxis angle was calculated between the net displacement and the direction toward the source. The signed chemotaxis index was calculated by projecting the net displacement onto the direction toward the chemical source and dividing this projected displacement by the total trajectory length [20, 21]. Positive values therefore indicate net movement toward the source, negative values indicate net movement away from the source, and values close to zero indicate little net movement along the source direction. Chemotaxis angles and indices are reported as medians with interquartile ranges, and indices additionally as means ± standard deviations. The complete analysis script, *Achromatium*_chemotaxis_index_analysis.py, is provided as Supplementary Code 1.

### Visualization of single cell trajectories

For all microscopic analyses, time-lapse images were recorded at regular intervals ranging from 10 to 60 seconds during the experiment. The individual frames were overlaid using the software *Picolay* (www.picolay.de) (Cypionka, Völcker, & Rohde, 2016) to reconstruct the trajectories of individual cells. To visualize the directionality of the movement, single cell traces (as shown in Figure 2) were generated from the same time-lapse images and color-coded using the software *Trace* (www.microbial-world.com/freewarelist).

### Calculation of single cell velocities

Single cell velocities were calculated from time-lapse images based on the distance traveled by each cell over several exposures and the time elapsed between these images.

Single cell velocities were calculated for cells moving along gradients of oxygen (n=15), nitrate (n=16), and sulfide (n=15). In addition, velocities were determined for cells exposed to different oxygen concentrations, including anoxic (n=15), hypoxic (n=20), and oxic (n=27) conditions. All determined velocities are listed in Table S4.

### Statistical analyses

Statistical analyses were performed in PAST (v. 5.3)[61] The two datasets obtained under uniform oxic conditions were combined into one oxic group (n = 27). Homogeneity of variance was evaluated using a median-centered Levene test. Because variances differed significantly among conditions (F(5,102) = 7.60, p = 4.21 × 10^−6^), single-cell velocities were compared using a Kruskal–Wallis test with tie correction, followed by Dunn’s pairwise post hoc tests with Bonferroni correction. All tests were two-sided, with α = 0.05.

### Bioinformatic analysis

All public genomes of *Achromatium* spp. [3, 4, 6, 7] were annotated de-novo using Bakta (V1.11.4; [62]). Additionally, all genes called during annotation were analyzed using the command line interface of InterProScan (V.5.77-108 [63]) and all available databases for protein and domain annotation.

Chemotaxis receptors (methyl-accepting chemotaxis proteins, MCPs) were identified and classified based on conserved domain architecture derived from InterProScan annotations. Proteins were considered bona fide MCPs only if they contained a canonical methyl-accepting (MA) signaling domain (Pfam PF00015), with optional presence of HAMP linker domains (PF00672). Proteins lacking PF00015 were excluded from further analysis. MCPs were subsequently subclassified according to the presence of specific sensory domains detected within the same polypeptide. BLUF-domain MCPs were defined by the presence of Pfam PF04940 and the corresponding conserved signatures SM01034, PS50925, and InterPro IPR007024. LOV-domain MCPs were identified by Pfam PF03441 and the keywords “LOV, or light-oxygen-voltage”. Photoactive yellow protein (PYP) MCPs were identified by Pfam PF00818, Super Family SSF55785 and the keywords “PYP, or photoactive yellow protein.” Aer-type redox-sensing MCPs were defined by the presence of the FAD-binding PAS domain Pfam PF00989 and PF13426. PilJ-type chemoreceptors were identified using Pfam PF13675 and InterPro IPR029095. Assignment to a sensory class required co-occurrence of the respective sensory domain with the MCP signaling domain (PF00015) within a single protein sequence. Proteins containing more than one sensory domain were annotated as multi-sensory MCPs, whereas MCPs lacking any of the defined sensory Pfam matches were classified as uncharacterized. This domain-based classification ensured reproducible separation of MCP subclasses independent of gene name annotation. The python script used to identify and classify MCPs based on the InterProScan annotation is given in the supplementary files. For non-canonical MCPs, the protein name was extracted from the MCP annotation and all the domains of these proteins were extracted from the InterProScan annotation, and analyzed manually.

*Achromatium* transcriptomes [3] were mapped against the above called genes using BBMap to obtain an overall qualitative picture on which genes were expressed.

### Diffusion model

A two-dimensional diffusion model was implemented to estimate the spatial distribution of dissolved H_2_S, O_2_^−^, NO_3_^−^, and NO_2_^−^ in a 16.5 × 16.5 mm diffusion chamber over time (10 and 60 min). The computational domain was discretized on a uniform 250 × 250 grid, corresponding to a spatial resolution of 0.064 mm. Transport was described by Fick’s second law, *∂C*/*∂t* = *D∇*^2^*C*, and solved using an explicit finite-difference scheme with a five-point central-difference approximation of the Laplacian. For the simulations, diffusion coefficients of 1.6 × 10^−3^ mm^2^ s^−1^ for H_2_S, 1.5 × 10^−3^ mm^2^ s^−1^ for NO_3_^−^, 1.5 × 10^−3^ mm^2^ s^−1^ for NO_2_^−^, and 1.8 × 10^−3^ mm^2^ s^−1^ for O_2_ were used. The initial concentration of H_2_S, NO_3_^−^, and NO_2_^−^, in the source region were set according to the different experiments, For O_2_, 0.27 mM were used to reflecting air/water equilibrium. The initial concentration elsewhere in the chamber was set to zero. The source occupied a 2 × 1 mm region adjacent to one chamber corner. A 2 × 0.1 mm opening at the lower edge of this source region was represented as a zero-concentration boundary (sink) at every time step, representing loss through the chamber opening. The integration time step was selected according to the explicit-scheme stability condition, *dt* = 0.495 *dx*^2^ /*D*, and the number of time steps was chosen to represent 600 s of diffusion. Concentration fields were displayed on a logarithmic scale from 0.001 to 100 mM; values below 0.001 mM were clipped only for visualization and not for the numerical calculation. The model describes diffusion only and does not include advection, chemical transformation, H_2_S oxidation, adsorption, biological uptake, or changes in diffusion coefficient caused by temperature, viscosity, or the presence of agar. The model was implemented in Python.

## Data availability

No new sequence data was generated in this study. The results of sequence data analyses of public *Achromatium* data were deposited on Zenodo at: https://doi.org/10.64898/2026.03.30.715255

## Acknowledgements

We are grateful to Hans-Dietrich Babenzien, who started the studies on *Achromatium* at Lake Stechlin and inspired us to continue his work after his retirement. Also, we kindly acknowledge Jana Schmidt and Frank Meyerjürgens for their technical assistance and the Leibniz Association for financial support to HPG. DI and the molecular work were funded through DFG grant “Eco-evolution and cellular functionality of the heterozygous bacterial genus *Achromatium*” (project number 465407921) awarded to DI.

