## Supplementary material for "Chemotaxis and motility of *Achromatium oxaliferum* in response to oxygen, sulfide, and nitrate (*)"

**Supplementary files**

**Video S1: Time-lapse video of a dividing *Achromatium* cell at 20× speed, illustrating its rolling and flipping movements.**

**Supplementary figures**

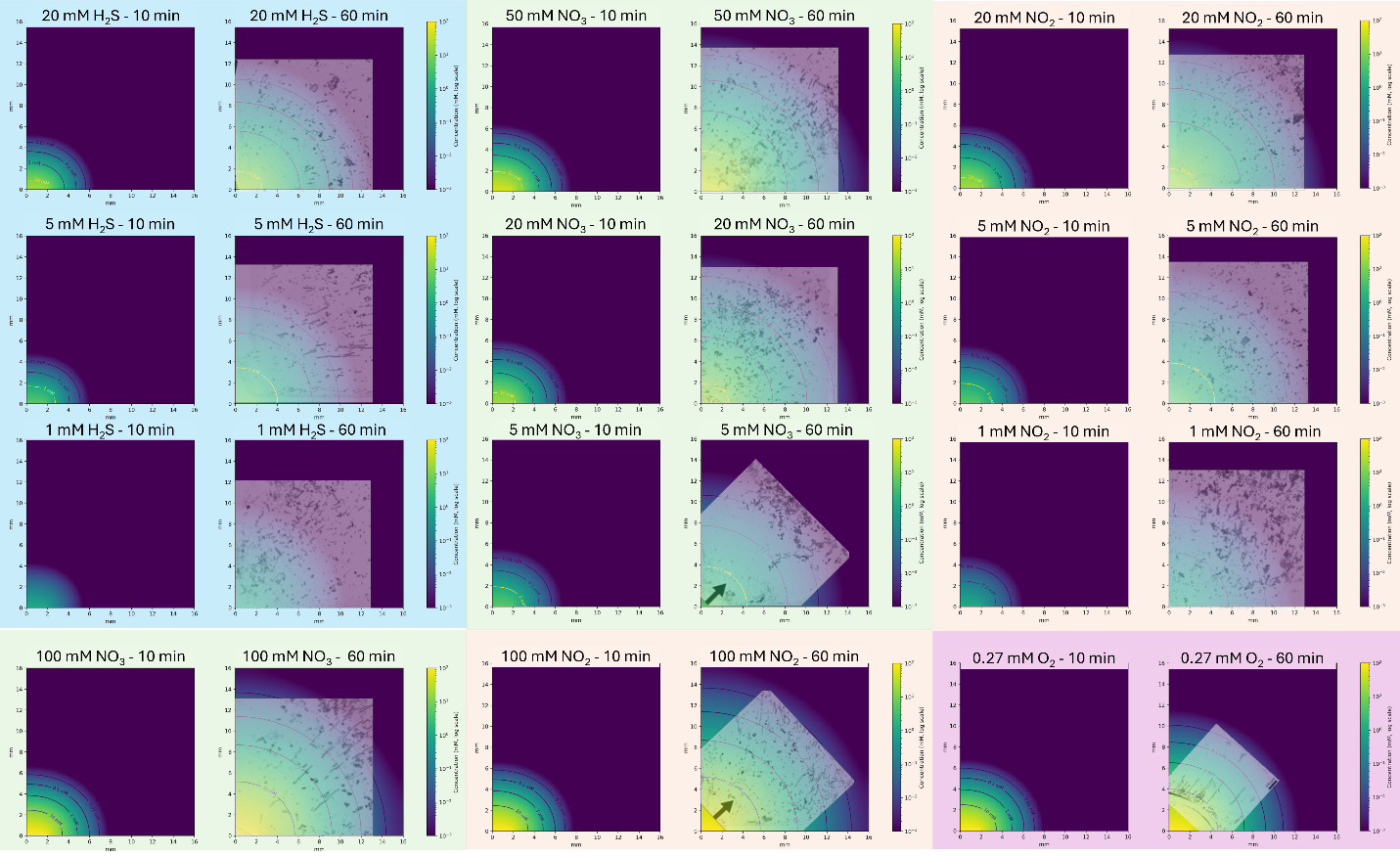

**Figure S1.** Diffusion models of individual solutes after 10 and 60 minutes of chemotaxis experiments. Agar plugs were soaked in solute solutions unitl saturation and placed at the opening at the lower left corner (2 mm opening). Photomicrographs of 60 minuntes overlaid chemotaxis assays were superimposed on the matching model.

**Supplementary tables**

**Table S1:** Chemotactic response of *Achromatium* cells after exposure to nitrate, nitrite, and sulfide at different concentrations. Reported is the fraction of cells exhibiting a chemotaxis response at a given solute concentration in percent (%).

| **Experiment** | **Concentration** | **Number of motile cells** | **Number of cells performing chemotaxis** | **Percentage of cells performing chemotaxis** |
| --- | --- | --- | --- | --- |
| Sulfide | 1 mM | 105 | 38 | 36.2 |
| Sulfide | 5 mM | 219 | 70 | 32.0 |
| Sulfide | 20 mM | 103 | 23 | 22.3 |
| Nitrite | 1 mM | 431 | 60 | 13.9 |
| Nitrite | 5 mM | 149 | 15 | 10.1 |
| Nitrite | 20 mM | 62 | 18 | 29.0 |
| Nitrite | 100 mM | 79 | 40 | 50.6 |
| Nitrate | 5 mM | 119 | 0 | 0.0 |
| Nitrate | 20 mM | 140 | 0 | 0.0 |
| Nitrate | 50 mM | 116 | 30 | 25.9 |
| Nitrate | 100 mM | 146 | 38 | 26.0 |

**Table S2**: Diversity of MCPs.

| **Complexity Level** | **Sensory Domains** | **Light Sensor** | **Notes** |
| --- | --- | --- | --- |
| Simple Canonical | Four helix bundle (MCP core sensory); PAS (single) | No | Basic attractant/repellent sensors; HAMP → adaptation. |
| Moderate Canonical | Cache (1-2) + PAS | No | Nutrient/chemical binding; Cache for periplasmic ligands. |
| Complex Canonical | Multi-PAS/Cache + PYP-like (SSF55785) | Yes (PYP/PAS) | Photoreceptor MCPs; pas2/PYP for light chemotaxis. |
| Non-Light Variants | CHASE3; Double sensory/PAS | No | Amino acid/peptide sensors; no BLUF detected. |

**Table S3**: Diversity of light sensors.

| **Sensor type** | **Light Sensor Domain(s)** | **Effector/Output** | **Notes** |
| --- | --- | --- | --- |
| Simple sensory HK | N-PAS/PYP-like (SSF55785) | HK (HisKA + HATPase_c) → Direct His→RR | Standard two-component system (TCS) for phototaxis/chemotaxis. |
| Hybrid HK | N-PAS/PYP → HK → C-terminal REC | REC (PF00072, CheY-like); Internal His→Asp relay | Unidirectional relay within hybrid kinase. |
| Hybrid + Hpt | N-PAS/PYP → HK → 1–2 REC → C-Hpt | Hpt (PF01627, SSF47226); Multi-step His→Asp→His→RR | Phosphotransfer relay for distributed signaling. |
| Complex/multi-relay | N-CHASE/GAF + multi-PAS/PYP → HK → 2 REC → Hpt (± fusions) | Extended relay + chemotaxis (e.g., methyltransferase) | Advanced adaptation |
| GGDEF-only | PYP-like PAS (SSF55785) or PAS | GGDEF cyclase (PF00990); c-di-GMP synthesis | Light-regulated motility; response regulator coupled. |
| EAL-only | PYP-like PAS or BLUF (PF04940) | EAL phosphodiesterase (PF00563); c-di-GMP degradation | Degradation-focused control. |
| Dual GGDEF+EAL | PYP-like PAS, PAS, BLUF | GGDEF (PF00990) + EAL (PF00563) | Degenerate allosteric regulation of c-di-GMP turnover. |
| Other | BLUF (PF04940), PAS/PYP-like | Phosphatase (PP2C, PF13672); standalone |  |
