## Supplementary material for "Chemotaxis and motility of *Achromatium oxaliferum* in response to oxygen, sulfide, and nitrate (*)": Table S4

**Table S4: Parameters and motility characteristics of Achromatium cells recorded during chemotaxis and motility assays. The table summarizes the imaging parameters, duration of continuous movement, distance travelled, and calculated cell velocities for each experiment.**

| Experiment | Number of cell | Time between frames (min) | Total number of frames recorded | Convert pixel → µm | Period of continous movement (frames) | Period of continous movement (pixel) | Period of continous movement (s) | Period of continous movement (µm) | Velocity ($\mu m\ s^{-1}$) | Median velocity ($\mu m\ s^{-1}$) |
|---|---|---|---|---|---|---|---|---|---|---|
| Oxygen chemotaxis (O2 content of air) | 1 | 0.50 | 60.00 | 0.06 | 5.00 | 17.00 | 300.00 | 294.67 | 1.96 | 1.57 |
| Oxygen chemotaxis (O2 content of air) | 2 | 0.50 | 60.00 | 0.06 | 3.00 | 19.00 | 180.00 | 329.33 | 3.66 | |
| Oxygen chemotaxis (O2 content of air) | 3 | 0.50 | 60.00 | 0.06 | 8.00 | 14.00 | 480.00 | 242.67 | 1.01 | |
| Oxygen chemotaxis (O2 content of air) | 4 | 0.50 | 60.00 | 0.06 | 5.00 | 10.00 | 300.00 | 173.33 | 1.16 | |
| Oxygen chemotaxis (O2 content of air) | 5 | 0.50 | 60.00 | 0.06 | 8.00 | 14.00 | 480.00 | 242.67 | 1.01 | |
| Oxygen chemotaxis (O2 content of air) | 6 | 0.50 | 60.00 | 0.06 | 7.00 | 19.00 | 420.00 | 329.33 | 1.57 | |
| Oxygen chemotaxis (O2 content of air) | 7 | 0.50 | 60.00 | 0.06 | 5.00 | 17.00 | 300.00 | 294.67 | 1.96 | |
| Oxygen chemotaxis (O2 content of air) | 8 | 0.50 | 60.00 | 0.06 | 5.00 | 32.00 | 300.00 | 554.67 | 3.70 | |
| Oxygen chemotaxis (O2 content of air) | 9 | 0.50 | 60.00 | 0.06 | 6.00 | 18.00 | 360.00 | 312.00 | 1.73 | |
| Oxygen chemotaxis (O2 content of air) | 10 | 0.50 | 60.00 | 0.06 | 8.00 | 28.00 | 480.00 | 485.33 | 2.02 | |
| Oxygen chemotaxis (O2 content of air) | 11 | 0.50 | 60.00 | 0.06 | 10.00 | 31.00 | 600.00 | 537.33 | 1.79 | |
| Oxygen chemotaxis (O2 content of air) | 12 | 0.50 | 60.00 | 0.06 | 11.00 | 7.00 | 660.00 | 121.33 | 0.37 | |
| Oxygen chemotaxis (O2 content of air) | 13 | 0.50 | 60.00 | 0.06 | 9.00 | 12.00 | 540.00 | 208.00 | 0.77 | |
| Oxygen chemotaxis (O2 content of air) | 14 | 0.50 | 60.00 | 0.06 | 10.00 | 9.00 | 600.00 | 156.00 | 0.52 | |
| Oxygen chemotaxis (O2 content of air) | 15 | 0.50 | 60.00 | 0.06 | 18.00 | 11.00 | 1080.00 | 190.67 | 0.35 | |
| Nitrate chemotaxis (50 mM) | 1 | 1.00 | 60.00 | 0.70 | 5.00 | 95.00 | 300.00 | 136.69 | 0.46 | 0.37 |
| Nitrate chemotaxis (50 mM) | 2 | 1.00 | 60.00 | 0.70 | 4.00 | 67.00 | 240.00 | 96.40 | 0.40 | |
| Nitrate chemotaxis (50 mM) | 3 | 1.00 | 60.00 | 0.70 | 5.00 | 74.00 | 300.00 | 106.47 | 0.35 | |
| Nitrate chemotaxis (50 mM) | 4 | 1.00 | 60.00 | 0.70 | 5.00 | 91.00 | 300.00 | 130.94 | 0.44 | |
| Nitrate chemotaxis (50 mM) | 5 | 1.00 | 60.00 | 0.70 | 5.00 | 65.00 | 300.00 | 93.53 | 0.31 | |
| Nitrate chemotaxis (100 mM) | 1 | 1.00 | 60.00 | 0.70 | 12.00 | 174.00 | 720.00 | 250.36 | 0.35 | |
| Nitrate chemotaxis (100 mM) | 2 | 1.00 | 60.00 | 0.70 | 7.00 | 110.00 | 420.00 | 158.27 | 0.38 | |
| Nitrate chemotaxis (100 mM) | 3 | 1.00 | 60.00 | 0.70 | 5.00 | 87.00 | 300.00 | 125.18 | 0.42 | |
| Nitrate chemotaxis (100 mM) | 4 | 1.00 | 60.00 | 0.70 | 6.00 | 96.00 | 360.00 | 138.13 | 0.38 | |
| Nitrate chemotaxis (100 mM) | 5 | 1.00 | 60.00 | 0.70 | 7.00 | 115.00 | 420.00 | 165.47 | 0.39 | |
| Nitrate chemotaxis (100 mM) | 6 | 1.00 | 60.00 | 0.70 | 6.00 | 95.00 | 360.00 | 136.69 | 0.38 | |
| Nitrate chemotaxis (100 mM) | 7 | 1.00 | 60.00 | 0.70 | 8.00 | 123.00 | 480.00 | 176.98 | 0.37 | |
| Nitrate chemotaxis (100 mM) | 8 | 1.00 | 60.00 | 0.70 | 8.00 | 74.00 | 480.00 | 106.47 | 0.22 | |
| Nitrate chemotaxis (100 mM) | 9 | 1.00 | 60.00 | 0.70 | 5.00 | 73.00 | 300.00 | 105.04 | 0.35 | |
| Nitrate chemotaxis (100 mM) | 10 | 1.00 | 60.00 | 0.70 | 8.00 | 60.00 | 480.00 | 86.33 | 0.18 | |
| Nitrate chemotaxis (100 mM) | 11 | 1.00 | 60.00 | 0.70 | 5.00 | 71.00 | 300.00 | 102.16 | 0.34 | |
| Sulfide chemotaxis (5 mM) | 1 | 1.00 | 60.00 | 0.70 | 11.00 | 70.00 | 660.00 | 100.72 | 0.15 | 0.37 |

| | | | | | | | | | | |
|---|---|---|---|---|---|---|---|---|---|---|
| Sulfide chemotaxis (5 mM) | 2 | 1.00 | 60.00 | 0.70 | 5.00 | 79.00 | 300.00 | 113.67 | 0.38 | |
| Sulfide chemotaxis (5 mM) | 3 | 1.00 | 60.00 | 0.70 | 4.00 | 65.00 | 240.00 | 93.53 | 0.39 | |
| Sulfide chemotaxis (5 mM) | 4 | 1.00 | 60.00 | 0.70 | 5.00 | 90.00 | 300.00 | 129.50 | 0.43 | |
| Sulfide chemotaxis (5 mM) | 5 | 1.00 | 60.00 | 0.70 | 4.00 | 52.00 | 240.00 | 74.82 | 0.31 | |
| Sulfide chemotaxis (5 mM) | 6 | 1.00 | 60.00 | 0.70 | 4.00 | 66.00 | 240.00 | 94.96 | 0.40 | |
| Sulfide chemotaxis (5 mM) | 7 | 1.00 | 60.00 | 0.70 | 4.00 | 62.00 | 240.00 | 89.21 | 0.37 | |
| Sulfide chemotaxis (5 mM) | 8 | 1.00 | 60.00 | 0.70 | 5.00 | 99.00 | 300.00 | 142.45 | 0.47 | |
| Sulfide chemotaxis (5 mM) | 9 | 1.00 | 60.00 | 0.70 | 7.00 | 66.00 | 420.00 | 94.96 | 0.23 | |
| Sulfide chemotaxis (5 mM) | 10 | 1.00 | 60.00 | 0.70 | 4.00 | 67.00 | 240.00 | 96.40 | 0.40 | |
| Sulfide chemotaxis (5 mM) | 11 | 1.00 | 60.00 | 0.70 | 7.00 | 53.00 | 420.00 | 76.26 | 0.18 | |
| Sulfide chemotaxis (5 mM) | 12 | 1.00 | 60.00 | 0.70 | 5.00 | 51.00 | 300.00 | 73.38 | 0.24 | |
| Sulfide chemotaxis (5 mM) | 13 | 1.00 | 60.00 | 0.70 | 4.00 | 65.00 | 240.00 | 93.53 | 0.39 | |
| Sulfide chemotaxis (5 mM) | 14 | 1.00 | 60.00 | 0.70 | 4.00 | 60.00 | 240.00 | 86.33 | 0.36 | |
| Sulfide chemotaxis (5 mM) | 15 | 1.00 | 60.00 | 0.70 | 4.00 | 45.00 | 240.00 | 64.75 | 0.27 | |
| Chemotaxis experiments (anoxic control) | 1 | 0.50 | 60.00 | 0.70 | 7.00 | 91.00 | 420.00 | 130.94 | 0.62 | 0.44 |
| Chemotaxis experiments (anoxic control) | 2 | 0.50 | 60.00 | 0.70 | 6.00 | 70.00 | 360.00 | 100.72 | 0.56 | |
| Chemotaxis experiments (anoxic control) | 3 | 0.50 | 60.00 | 0.70 | 4.00 | 54.00 | 240.00 | 77.70 | 0.65 | |
| Chemotaxis experiments (anoxic control) | 4 | 0.50 | 60.00 | 0.70 | 5.00 | 45.00 | 300.00 | 64.75 | 0.43 | |
| Chemotaxis experiments (anoxic control) | 5 | 0.50 | 60.00 | 0.70 | 3.00 | 49.00 | 180.00 | 70.50 | 0.78 | |
| Chemotaxis experiments (anoxic control) | 6 | 0.50 | 60.00 | 0.70 | 6.00 | 71.00 | 360.00 | 102.16 | 0.57 | |
| Chemotaxis experiments (anoxic control) | 7 | 0.50 | 60.00 | 0.70 | 8.00 | 48.00 | 480.00 | 69.06 | 0.29 | |
| Chemotaxis experiments (anoxic control) | 8 | 0.50 | 60.00 | 0.70 | 8.00 | 74.00 | 480.00 | 106.47 | 0.44 | |
| Chemotaxis experiments (anoxic control) | 9 | 0.50 | 60.00 | 0.70 | 6.00 | 51.00 | 360.00 | 73.38 | 0.41 | |
| Chemotaxis experiments (anoxic control) | 10 | 0.50 | 60.00 | 0.70 | 10.00 | 62.00 | 600.00 | 89.21 | 0.30 | |
| Chemotaxis experiments (anoxic control) | 11 | 0.50 | 60.00 | 0.70 | 7.00 | 63.00 | 420.00 | 90.65 | 0.43 | |
| Chemotaxis experiments (anoxic control) | 12 | 0.50 | 60.00 | 0.70 | 4.00 | 47.00 | 240.00 | 67.63 | 0.56 | |
| Chemotaxis experiments (anoxic control) | 13 | 0.50 | 60.00 | 0.70 | 13.00 | 74.00 | 780.00 | 106.47 | 0.27 | |
| Chemotaxis experiments (anoxic control) | 14 | 0.50 | 60.00 | 0.70 | 7.00 | 41.00 | 420.00 | 58.99 | 0.28 | |
| Chemotaxis experiments (anoxic control) | 15 | 0.50 | 60.00 | 0.70 | 4.00 | 72.00 | 240.00 | 103.60 | 0.86 | |
| Chemotaxis experiments (oxic control) | 1 | 0.50 | 60.00 | 0.06 | 12.00 | 14.00 | 720.00 | 242.67 | 0.67 | 0.91 |
| Chemotaxis experiments (oxic control) | 2 | 0.50 | 60.00 | 0.06 | 7.00 | 43.00 | 420.00 | 745.33 | 3.55 | |
| Chemotaxis experiments (oxic control) | 3 | 0.50 | 60.00 | 0.06 | 7.00 | 14.00 | 420.00 | 242.67 | 1.16 | |
| Chemotaxis experiments (oxic control) | 4 | 0.50 | 60.00 | 0.06 | 17.00 | 16.00 | 1020.00 | 277.33 | 0.54 | |
| Chemotaxis experiments (oxic control) | 5 | 0.50 | 60.00 | 0.06 | 16.00 | 16.00 | 960.00 | 277.33 | 0.58 | |
| Chemotaxis experiments (oxic control) | 6 | 0.50 | 60.00 | 0.06 | 15.00 | 13.00 | 900.00 | 225.33 | 0.50 | |
| Chemotaxis experiments (oxic control) | 7 | 0.50 | 60.00 | 0.06 | 10.00 | 10.00 | 600.00 | 173.33 | 0.58 | |
| Chemotaxis experiments (oxic control) | 8 | 0.50 | 60.00 | 0.06 | 7.00 | 11.00 | 420.00 | 190.67 | 0.91 | |
| Chemotaxis experiments (oxic control) | 9 | 0.50 | 60.00 | 0.06 | 7.00 | 16.00 | 420.00 | 277.33 | 1.32 | |
| Chemotaxis experiments (oxic control) | 10 | 0.50 | 60.00 | 0.06 | 17.00 | 31.00 | 1020.00 | 537.33 | 1.05 | |
| Chemotaxis experiments (oxic control) | 1 | 0.50 | 60.00 | 0.70 | 12.00 | 108.00 | 720.00 | 155.40 | 0.43 | |
| Chemotaxis experiments (oxic control) | 2 | 0.50 | 60.00 | 0.70 | 6.00 | 40.00 | 360.00 | 57.55 | 0.32 | |
| Chemotaxis experiments (oxic control) | 3 | 0.50 | 60.00 | 0.70 | 8.00 | 129.00 | 480.00 | 185.61 | 0.77 | |
| Chemotaxis experiments (oxic control) | 4 | 0.50 | 60.00 | 0.70 | 7.00 | 98.00 | 420.00 | 141.01 | 0.67 | |
| Chemotaxis experiments (oxic control) | 5 | 0.50 | 60.00 | 0.70 | 7.00 | 112.00 | 420.00 | 161.15 | 0.77 | |
| Chemotaxis experiments (oxic control) | 6 | 0.50 | 60.00 | 0.70 | 8.00 | 129.00 | 480.00 | 185.61 | 0.77 | |

| | | | | | | | | | | |
|---|---|---|---|---|---|---|---|---|---|---|
| Chemotaxis experiments (oxic control) | 7 | 0.50 | 60.00 | 0.70 | 4.00 | 98.00 | 240.00 | 141.01 | 1.18 | |
| Motility (oxic conditions) | 1 | 0.50 | 40.00 | 0.63 | 4.00 | 90.00 | 240.00 | 144.00 | 1.20 | |
| Motility (oxic conditions) | 2 | 0.50 | 40.00 | 0.63 | 3.00 | 80.00 | 180.00 | 128.00 | 1.42 | |
| Motility (oxic conditions) | 3 | 0.50 | 40.00 | 0.63 | 5.00 | 89.00 | 300.00 | 142.40 | 0.95 | |
| Motility (oxic conditions) | 4 | 0.50 | 40.00 | 0.63 | 3.00 | 64.00 | 180.00 | 102.40 | 1.14 | |
| Motility (oxic conditions) | 5 | 0.50 | 40.00 | 0.63 | 9.00 | 67.00 | 540.00 | 107.20 | 0.40 | |
| Motility (oxic conditions) | 6 | 0.50 | 40.00 | 0.63 | 3.00 | 119.00 | 180.00 | 190.40 | 2.12 | |
| Motility (oxic conditions) | 7 | 0.50 | 40.00 | 0.63 | 3.00 | 107.00 | 180.00 | 171.20 | 1.90 | |
| Motility (oxic conditions) | 8 | 0.50 | 40.00 | 0.63 | 3.00 | 90.00 | 180.00 | 144.00 | 1.60 | |
| Motility (oxic conditions) | 9 | 0.50 | 40.00 | 0.63 | 8.00 | 135.00 | 480.00 | 216.00 | 0.90 | |
| Motility (oxic conditions) | 10 | 0.50 | 40.00 | 0.63 | 4.00 | 76.00 | 240.00 | 121.60 | 1.01 | |
| Motility (hypoxic conditions) | 1 | 0.17 | 30.00 | 0.63 | 5.00 | 82.00 | 300.00 | 131.20 | 2.62 | 2.07 |
| Motility (hypoxic conditions) | 2 | 0.17 | 30.00 | 0.63 | 7.00 | 90.00 | 420.00 | 144.00 | 2.06 | |
| Motility (hypoxic conditions) | 3 | 0.17 | 30.00 | 0.63 | 6.00 | 81.00 | 360.00 | 129.60 | 2.16 | |
| Motility (hypoxic conditions) | 4 | 0.17 | 30.00 | 0.63 | 4.00 | 82.00 | 240.00 | 131.20 | 3.28 | |
| Motility (hypoxic conditions) | 5 | 0.17 | 30.00 | 0.63 | 6.00 | 71.00 | 360.00 | 113.60 | 1.89 | |
| Motility (hypoxic conditions) | 6 | 0.17 | 30.00 | 0.63 | 2.00 | 26.00 | 120.00 | 41.60 | 2.08 | |
| Motility (hypoxic conditions) | 7 | 0.17 | 30.00 | 0.63 | 3.00 | 44.00 | 180.00 | 70.40 | 2.35 | |
| Motility (hypoxic conditions) | 8 | 0.17 | 30.00 | 0.63 | 5.00 | 56.00 | 300.00 | 89.60 | 1.79 | |
| Motility (hypoxic conditions) | 9 | 0.17 | 30.00 | 0.63 | 4.00 | 54.00 | 240.00 | 86.40 | 2.16 | |
| Motility (hypoxic conditions) | 10 | 0.17 | 30.00 | 0.63 | 4.00 | 38.00 | 240.00 | 60.80 | 1.52 | |
| Motility (hypoxic conditions) | 1 | 0.17 | 30.00 | 0.63 | 6.00 | 73.00 | 360.00 | 116.80 | 1.95 | |
| Motility (hypoxic conditions) | 2 | 0.17 | 30.00 | 0.63 | 9.00 | 31.00 | 540.00 | 49.60 | 0.55 | |
| Motility (hypoxic conditions) | 3 | 0.17 | 30.00 | 0.63 | 6.00 | 41.00 | 360.00 | 65.60 | 1.09 | |
| Motility (hypoxic conditions) | 4 | 0.17 | 30.00 | 0.63 | 3.00 | 51.00 | 180.00 | 81.60 | 2.72 | |
| Motility (hypoxic conditions) | 5 | 0.17 | 30.00 | 0.63 | 7.00 | 59.00 | 420.00 | 94.40 | 1.35 | |
| Motility (hypoxic conditions) | 6 | 0.17 | 30.00 | 0.63 | 4.00 | 37.00 | 240.00 | 59.20 | 1.48 | |
| Motility (hypoxic conditions) | 7 | 0.17 | 30.00 | 0.63 | 4.00 | 66.00 | 240.00 | 105.60 | 2.64 | |
| Motility (hypoxic conditions) | 8 | 0.17 | 30.00 | 0.63 | 2.00 | 27.00 | 120.00 | 43.20 | 2.16 | |
| Motility (hypoxic conditions) | 9 | 0.17 | 30.00 | 0.63 | 5.00 | 68.00 | 300.00 | 108.80 | 2.18 | |
| Motility (hypoxic conditions) | 10 | 0.17 | 30.00 | 0.63 | 4.00 | 49.00 | 240.00 | 78.40 | 1.96 | |
