## Supplementary figures and images for "Chemotaxis and motility of *Achromatium oxaliferum* in response to oxygen, sulfide, and nitrate (*)"

### Supplementary Movie 1

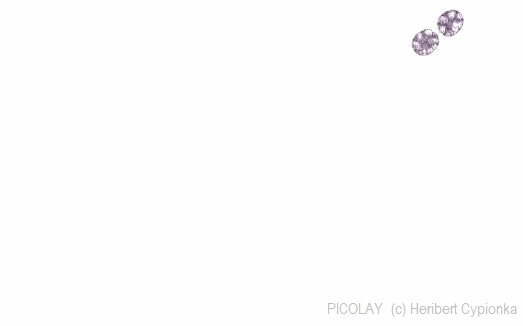
